# Complexity-stability relationship in empirical microbial ecosystems

**DOI:** 10.1101/2021.07.29.454345

**Authors:** Yogev Yonatan, Guy Amit, Jonathan Friedman, Amir Bashan

## Abstract

May’s stability theory [1, 2], which holds that large ecosystems can be stable up to a critical level of complexity, a product of the number of resident species and the intensity of their interactions, has been a central paradigm in theoretical ecology [3–7]. So far, however, empirically demonstrating this theory in real ecological systems has been a long-standing challenge, with inconsistent results [8]. Especially, it is unknown whether this theory is pertinent in the rich and complex communities of natural microbiomes, mainly due to the challenge of reliably reconstructing such large ecological interaction networks [9–11]. Here, we introduce a novel computational framework for estimating an ecosystem’s complexity without relying on a priori knowledge of its underlying interaction network. By applying this method to human-associated microbial communities from different body sites [12] and sponge-associated microbial communities from different geographical locations [13], we found that in both cases the communities display a pronounced trade-off between the number of species and their effective connectance. These results suggest that natural microbiomes are shaped by stability constraints, which limit their complexity.

---

Robert May has theoretically established that the stability of a large ecosystem of *N* randomly interacting species is determined by the complexity of the community matrix *M*, which describes the inter-species effects around an equilibrium point [1, 2, 14]. The complexity is defined as *α*^2^*NC*, where *α*^2^ and *C* are the variance and the density (‘Connectance’) of the non-zero off-diagonal elements of *M*, respectively. When the diagonal elements are scaled to −1, the ecosystem can be stable as long as it satisfies the stability criterion 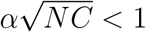.

This theory has been confirmed numerically for ecosystems with unspecified dynamics as well as Generalized Lotka-Volterra (GLV) dynamics with random inter-species interactions, and has been extended to various other types of interactions and network topologies [3–6, 15, 16]. May’s original qualitative principle remains robust: ecosystems with a large number of species are expected to be stable only when their other complexity parameters, i.e., the connectance *C* and the coupling strength *α*, are comparatively small [17], as illustrated in Fig. 1a.

**Figure 1:**
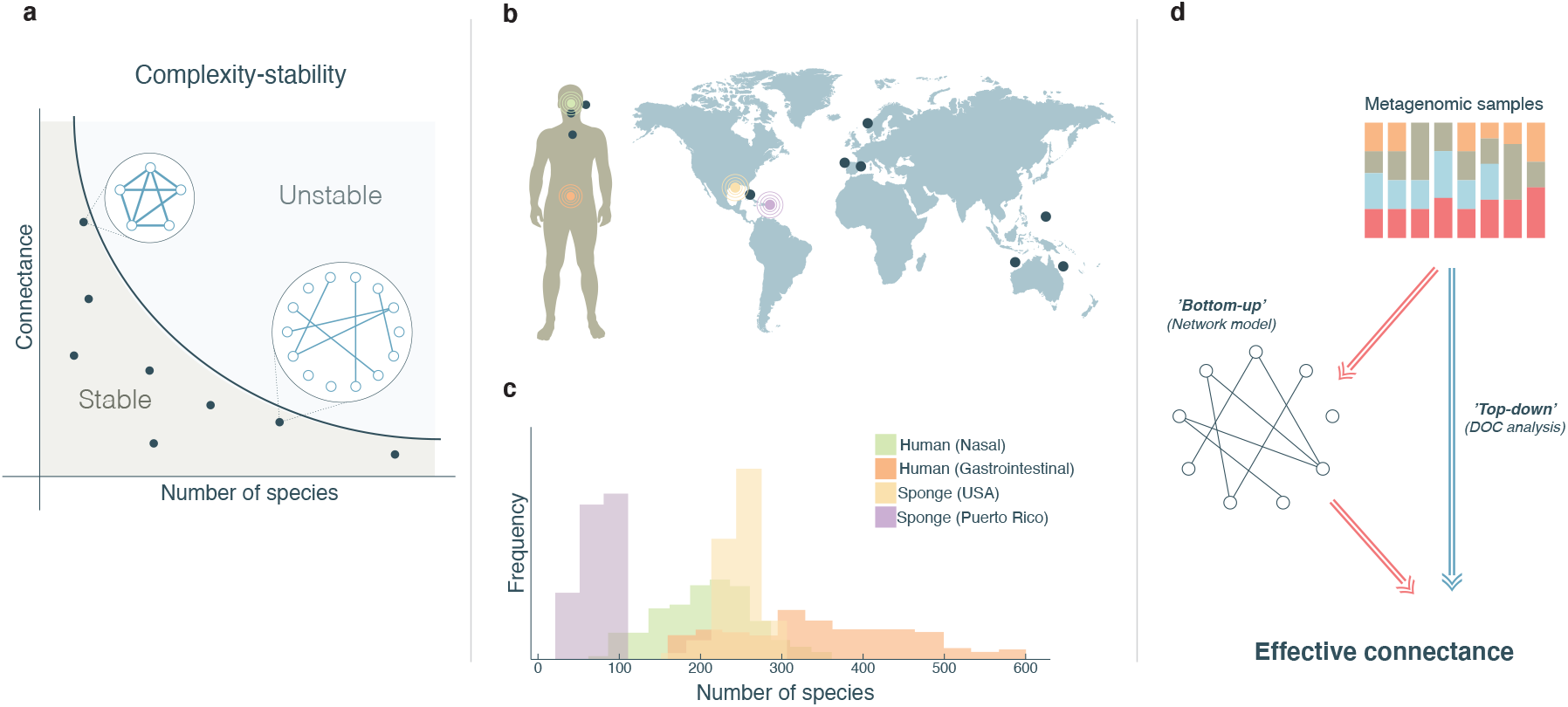
Observing complexity-stability patterns in natural microbial communities without network reconstruction. **a**, The complexity-stability paradigm predicts that there is an upper limit to the overall complexity of stable ecosystems, such that it would be improbable to observe communities with both a large number of species and a high degree of inter-species interactions (’Connectance’). While the number of species is, in principle, an observable measure, the networks of inter-species interactions are generally unknown. Dots in the connectance-number-of-species plane which are below the critical stability line represent viable ecosystems. **b**, In this study we analyze human-associated microbial samples from different body sites and sponge-associated microbial samples from diverse geographical locations, representing diverse ecological environments and species assemblages. **c**, The number of observed species (OTUs) in different samples from the same ecological environment typically distributed around a characteristic value, which vary substantially across the environments. **d**, A bottom-up approach for evaluating the connectance of an ecosystem from cross-sectional data relies on prior reconstruction of a detailed network model. Alternatively, the effective connectance can be directly extracted from the cross-sectional data without network inference, in a top-down approach, using the DOC method.

Empirical demonstration of May’s theory in real-world systems focuses on analysis of foodwebs, i.e., carefully constructed networks of ecological interactions between the species [18]. By comparing different food-webs, researchers investigated the relationship between the number of species, the connectance, and the community matrices’ stability parameters [19–25]. The conclusions, however, are inconsistent, leaving the question of whether May’s complexity-stability theory is indeed a governing principle in empirical ecosystems unanswered [8].

The recent surge in metagenomic surveys of natural microbial communities has opened a unique opportunity to test whether stability requirements constraint the complexity of real-world ecosystems in an unprecedented manner. First, large-scale projects have collected and processed high-throughput data from diverse environments [12, 26] using methodical protocols, such as 16S or whole-genome sequencing [27], with consistent ecological definitions, e.g., operational taxonomic units (OTUs) as a standardised definition of ‘species’. This resolves some of the inherent difficulties in previous comparisons of food-webs that stemmed from natural differences between field experiments, and from subjective definitions of species. Second, the number of resident species in a single microbial community is generally much larger than any other studied natural ecosystem, and can vary from a few dozens to thousands across different ecological environments. These unique features of microbial communities provide an ideal basis for performing an equitable test of May’s theory across ecosystems with a broad range of sizes.

Yet, assessing the complexity of inter-species interactions in the microbiome remains highly challenging. In the absence of a priori knowledge of the microbial ecological interaction networks, efforts to reconstruct them focus on inferring population dynamic models from temporal abundance data, i.e., the abundance time-series of each taxon in the microbial community [28–34]. This requires high-quality time series data and well-designed controlled experiments, which are currently scantily available, and rarely cover diverse ecological habitats [35–37]. The complexity of the community matrix cannot be extracted from co-occurrence networks either, since they do not encode any causal relations or direct ecological interactions [9, 38, 39].

To overcome these issues, we developed a novel computational framework for evaluating the system’s complexity from a cohort of cross-sectional samples, without inferring the complete interaction network. Instead, we directly evaluate the ‘effective connectance’ *α*^2^*C* of microbial ecosystems, which we define as the product of the interaction strength, *α*^2^ and the connectance, *C*, in a ‘top-down’ fashion, by testing how sensitive the abundances of the species are to the presence of additional species. Using this method we analyze the complexity-stability relationship in simulated GLV dynamics as well as in diverse natural microbial communities from human and marine environments.

## Evaluating the ‘effective connectance’ of complex ecosystems

We evaluate the effective connectance of the underlying dynamics, without inferring the detailed interaction network, by analyzing the relationship between two beta-diversity measures, the ‘dissimilarity’ and the ‘overlap’ of different sample pairs in a cohort. Consider two abundance profiles, **x** = (*x*_1_*, …, x_N_*) and **y** = (*y*_1_*, …, y_N_*), of *N* species. The overlap *Q*(**x**, **y**) is a measure of the similarity between the species assemblages of **x** and **y**, and the dissimilarity *R*(**x**, **y**) is a measure of the difference between the renormalized abundance profiles of the shared species, 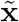 and 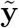 (see Methods and Supplementary Information Sec. 1.1.1). The dissimilarity and overlap values calculated for all sample pairs define the Dissimilarity-Overlap Curve (DOC), which has been previously used to test the existence of universal dynamics in human-associated microbial communities [40]. If the underlying dynamics that determine the abundance profiles are universal, the DOC will have a characteristic negative slope, 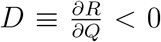, at the high overlap region, i.e., increased similarity between the abundance profiles of sample pairs with similar species assemblages.

Here, we show how the magnitude of the DOC’s slope can serve as a measure for the system’s effective connectance. In an ecosystem with denser and stronger connections between the species (higher *α*^2^*C*), even a small difference between the assemblages would lead to a large difference in their abundance profiles, and thus, a steeper DOC’s slope. The significance of *D* can be illustrated via the following hypothetical scenario. Consider two samples with similar species assemblages (large overlap) and similar abundance profiles (low dissimilarity). A minimal change in the overlap and dissimilarity between them can be applied by adding a new species to only one of them. While the overlap would decrease, the magnitude of change in the dissimilarity between the abundance profiles would depend on the effective connectance, i.e., how much the added species affects the abundances of the other species in its community. When calculated over all sample pairs, *D* describes the average effective connectance in a cohort.

The relation between *D* and *α*^2^*C* can be demonstrated analytically in the case of a GLV model of ecological dynamics, in which the underlying dynamics are described by *C* and *σ*, the probability (density) and characteristic strength of the inter-species interactions, respectively. We demonstrate in Supplementary Information Sec. 1.3 that the effective connectance *α*^2^*C*, which characterizes the resulting effects near an equilibrium point, is directly proportional to *σ*^2^*C*. Here, we show that the magnitude of the DOC’s slope, *D*, is proportional to *σ*^2^*C*, and hence can be used as a measure of *α*^2^*C*.

In the GVL case, the shape of the DOC at the high-overlap region, i.e., the relation between the dissimilarity *R* (measured using Euclidean distance) and the overlap *Q* is given by:

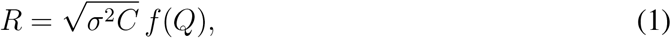

where 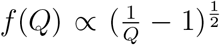 (see Supplementary Information Sec. 1.2). Thus, the magnitude of the DOC’s slope *D*, is 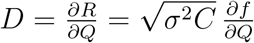, where 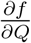 is relatively constant at 0.5 < *Q* < 0.9 (See Supplementary Information Fig. 1). Therefore, in this region,

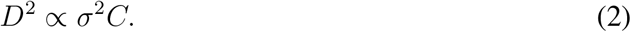

We suggest that for the GLV models, *σ*^2^*C* can be directly evaluated from a cohort of steady states, without the need to infer the complete interaction network.

To test the relation presented in Eq. (2), we simulate cohorts of steady states with different combinations of *C* and *σ* (see Methods). Subsequently, using the steady states, we evaluate *D* by a linear regression process (as demonstrated in Fig. 2a), and compare it with the predefined *σ*^2^*C* of each cohort. Figures 2a and 2b demonstrate that the DOC’s slope is steeper for denser (higher *C*) and stronger (higher *σ*) interaction networks. The squared DOC’s slope *D*^2^ is linearly related to both *C* and *σ*^2^ (Fig. 2c,d). The relation from Eq. (2) is demonstrated in Fig. 2e, exhibiting a clear linear relation between *σ*^2^*C* and *D*^2^. These results were found to be consistent under various scenarios, such as mutualistic, competitional, and predator-pray dynamics (See Supplementary Information Sec. 2). In conclusion, we demonstrate how the effective connectance can be recovered from cross-sectional data without any inference of the interaction network.

**Figure 2:**
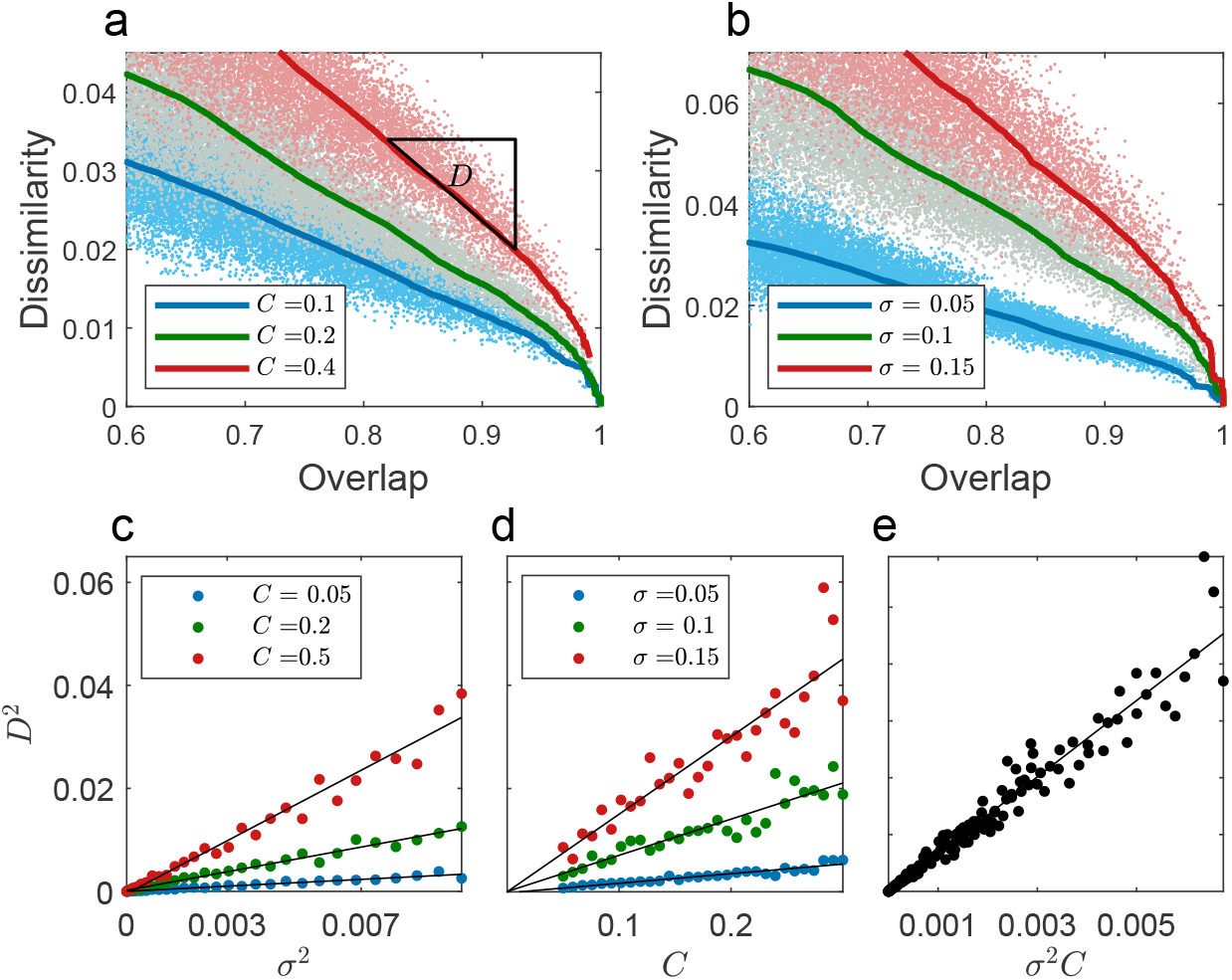
Dissimilarity-overlap analysis of cross-sectional data uncovers the predetermined connectance of the underlying GLV dynamics. **a**, Demonstration of the DOC’s slope for different connectance values, *C*. Three sets of *m* = 200 steady states are generated from different GLV models of *n* = 100 species with the same characteristic interaction strength (*σ* = 0.07) but different connectance values (*C* = 0.1, 0.2, 0.4). Points of the same color represent the overlap and dissimilarity (Euclidean distance) between all pairs of steady states from the same GLV model. The solid lines represent the corresponding DOCs (calculated as the moving average of the dissimilarity). The slope *D* for the case of *C* = 0.4 (red curve) is represented by the black triangle. **b**, DOC of three GLV models with the same connectance (*C* = 0.2), but with different interaction strengths (*σ* = 0.05, 0.1, 0.15). Both denser ecological networks (larger *C*) and stronger interactions (larger *σ*) lead to a steeper DOC. **c**, Relationship between *D*^2^ and the interactions’ characteristic strengths, *σ*^2^. Points of the same color exhibit linear relations between the *σ*^2^ and *D*^2^ values of GLV models with a fixed *C* (*R*^2^ *>* 0.93 in all three cases). For each GLV model, a set of *m* = 100 steady states was simulated, and the slope *D* was evaluated from a linear fit over the points with overlap values between 0.7 and 0.8. 30 models were generated for each *C* value. **d**, Relationship between *D*^2^, and the connectance, *C*. Points of the same color exhibit linear relations between *C* and *D*^2^ values of GLV models with fixed *σ* (*R*^2^ *>* 0.87 in all three cases). **e**, The *D*^2^ values of all former simulations presented in **c** and **d** are plotted with respect to the product *σ*^2^*C*. In this case, the product *σ*^2^*C* is linearly related to *D*^2^ as well (*R*^2^ = 0.95).

## The effective connectance of natural microbial communities

Next, we apply our method to metagenomic data of real microbial communities, and test whether they are constrained by the stability requirements. May’s stability theory predicts that it whould be improbable to observe real ecosystems with both a large number of species and high effective connectance. The required datasets for such analysis should consist of metagenomic samples from diverse ecological environments, where the number of samples from each environment should be sufficient for applying the DOC method. In addition, in order to minimize potential biases due to different experimental and processing procedures, we restrict ourselves to analyze only datasets generated as part of the same study.

We found two suitable studies that met these requirements, the Human Microbiome Project (HMP), investigating human-associated microbial communities from different body sites [12], and the Sponge Microbiome Project (SMP), which explores sponge-associated microbial communities from different geographical locations and host species [13]. In this work, we consider microbial species as OTUs. The microbial samples, which represent the abundance profiles of the OTUs in each local community, are divided into cohorts of similar ecological environments. We require at least 35 samples in each cohort to allow estimation of the DOC’s slope (see Supplementary Information Sec. 4). The primary patterns of variation in community structure across human-associated microbial samples are related to the body habitat group (oral, skin, gut, and airways) [12], thus, we divide them to cohorts according to the body site. Local communities of sponge-associated microbial samples collected from different hosts are divided according to the geographical sampling location, which is their main source of variation [41], followed by unsupervised clustering, in order to focus on cohorts that are as homogeneous as possible (see Methods). Finally, we analyze 13 human-associated cohorts and 15 sponge-associated cohorts of microbial samples.

For real metagenomic data, each cohort has its own characteristic range of overlap values. Therefore, we choose to calculate the DOC’s slopes over the 25% points with the highest overlap values in each cohort, i.e., the overlap fitting range is environment-specific (See Supplementary Information Sec. 3, validation using simulated data from GLV dynamics). In addition, in order to measure the dissimilarity between real ecological communities, we use the root Jensen-Shannon divergence (rJSD), a distance metric, which is less sensitive to biases caused by different numbers of species than the Euclidean distance (In Supplementary Information Sec. 5.2 we demonstrate that our results are qualitatively consistent for different dissimilarity measures). As shown in Fig. 3, the DOCs of all cohorts exhibit a clear negative slope, as expected in case of consistent (’universal’) dynamics [40], whereas the magnitudes of the slopes vary substantially across the different microbial communities. (Supplementary Fig. 12 shows that the variance among the slopes is observed even across cohorts of very similar overlap ranges, suggesting that the slope is an intrinsic feature of the cohort, and not an artifact stemming from the overlap range alone.)

**Figure 3:**
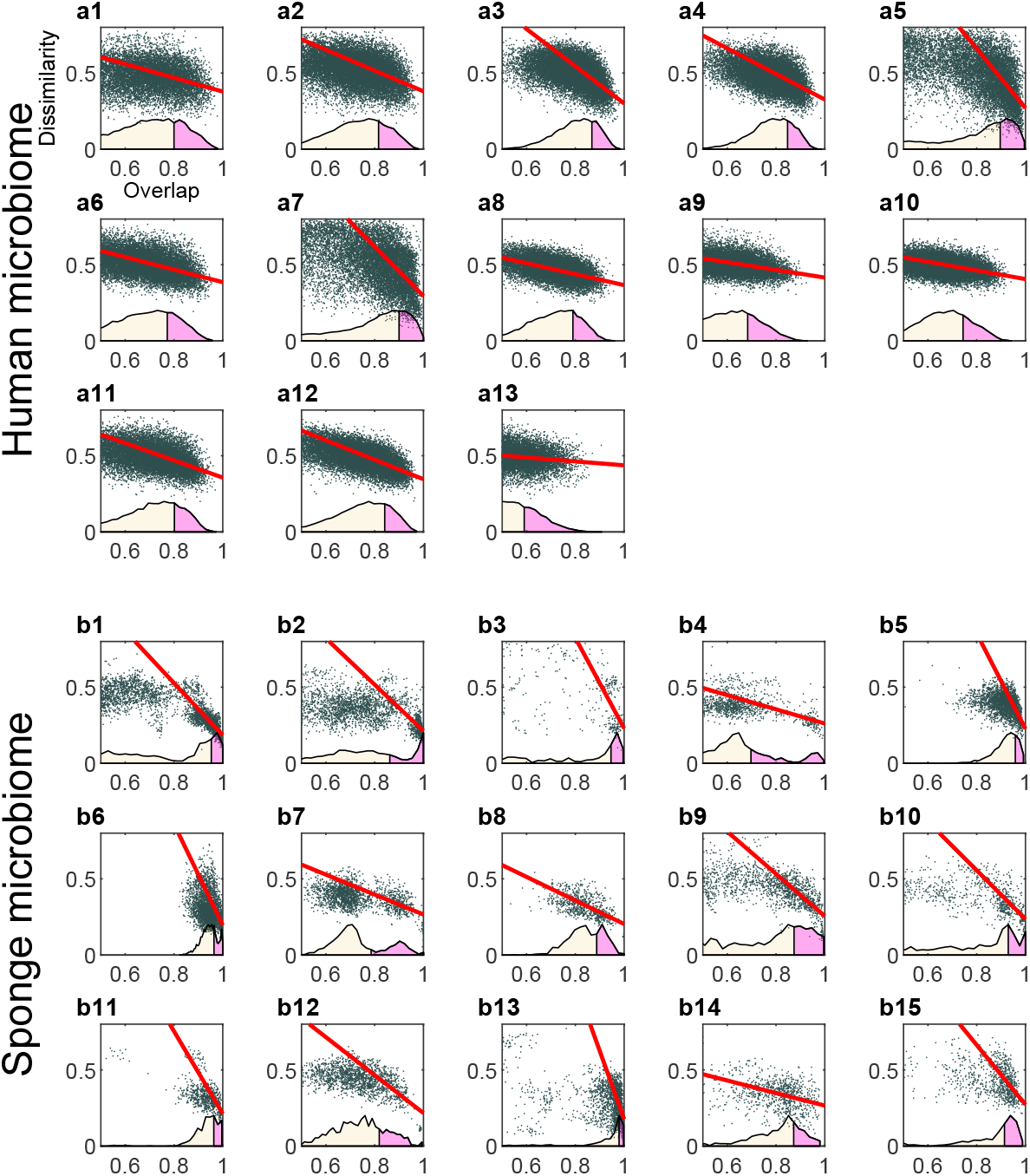
Dissimilarity-overlap curves across real-world microbial communities from diverse ecological environments. Human-associated microbial communities from the HMP: Anterior nares (**a1**), Attached keratinized nares (**a2**), Buccal mucosa (**a3**), Hard palate (**a4**), Left retroauricular crease (**a5**), Palatine tonsils (**a6**), Right retroauricular crease (**a7**), Saliva (**a8**), Subgingival plaque (**a9**), Supragingival plaque (**a10**), Throat (**a11**), Tongue dorsum (**a12**), Stool (**a13**). Sponge-associated microbial communities communities from the SMP: Australia (**b1**-**b2**), Puerto Rico (**b3**), Guam (**b4**), Norway (**b5**), Spain (**b6**-**b11**), USA (**b12**-**b15**). Dots represent the dissimilarity and overlap between all sample pairs. The overlap distributions of the sample pairs are depicted at the bottom, where the top 25% of each overlap distribution is marked in pink. The straight red lines are calculated using a linear fit over this overlap range, and are used to calculate the slopes of the DOCs. The diverse slopes of the DOCs observed both in the HMP and SMP studies suggest differences in their underlying effective connectance.

## Complexity-stability relationship in empirical microbial ecosystems

We investigate the complexity-stability relationship in real ecosystems by comparing the effective connectance, measured as the DOC slope, with the number of resident species. To verify our methodology, we apply it to simulated data generated from GLV models, which represent ecosystems formed under stability constraints, i.e., they are as large as possible, and as complex as possible whilst remaining stable (See Methods). As shown in Fig. 4a, the cohorts with high effective connectance (high *D*^2^) can be stable only when the number of species is relatively small, while a large number of species is found only in cohorts with low effective connectance, as expected from May’s stability criterion.

**Figure 4:**
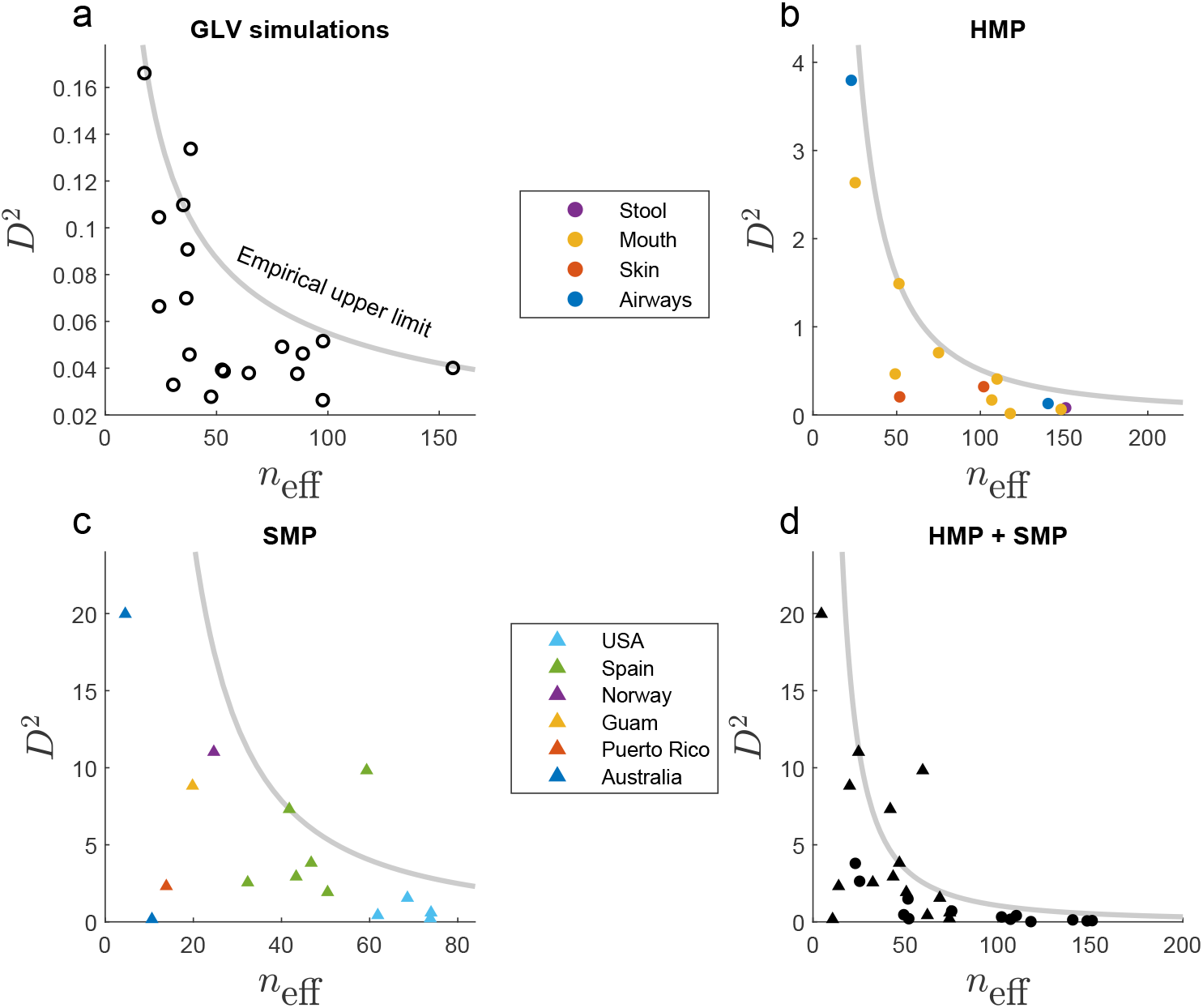
Complexity-stability patterns observed in real microbial ecosystems. **a**, The effective connectance, *D*^2^, versus the effective number of species, *n*_eff_, extracted from simulated cross-sectional data of GLV models that satisfy system stability (see Methods). Each circle represents a cohort of *m* = 50 alternative steady states of the same GLV model, where *D*^2^ is evaluated from the DOC (using the rJSD dissimilarity metric). The grey line represents the empirical upper limit of the data points with P-value = 1.1 × 10^−2^, using the randomly reshuffling test (see Methods), calculated from 10^4^ Monte-Carlo realizations. The number of cohorts is 20. The Pearson correlation rank is −0.528 (*P* = 0.016). Note that when using the Euclidean distance metric, the stability constraints of the GLV model are better represented by the empirical upper limit (See Supplementary Information Fig. 11). **b**, Dots represent cohorts of human-associated microbial samples, marked in different colors according to their body-site of origin. The number of samples in each cohort is larger than 145. The number of cohorts is 13. The P-value associated with the grey line is 10^−3^. The Pearson correlation rank is 0.75 (*P* = 0.003). **c**, Triangles represent cohorts of sponge-associated microbial samples. Different colors represent different geographical locations. The number of samples in each cohort is larger than 35. The number of cohorts is 15. The P-value associated with the grey line is 0.027. The Pearson correlation rank is −0.51 (*P* = 0.05). **d**, A plot of the data points from both **b** and **c** jointly exhibiting the complexity-stability pattern with a *P* < 10^−3^, associated with the grey line. The Pearson correlation rank is −0.52 (*P* = 0.003).

To apply this to real world microbial ecosystems, we must first estimate the number of resident species from genetic surveys. This is a challenging task, mainly due to the large number of low abundance species, which are difficult to detect. To account for that, we repeat our analysis using different, already established, ecological measures, and examine whether the general complexity-stability pattern is consistent with all of them. Specifically, we estimate the number of species in each microbial community using three alternative definitions: i) The number of observed OTUs, ii) The number of most abundant OTUs that sum up to 90% of the total abundance, and iii) The Shannon effective number of OTUs, (*n*_eff_) [39, 42], which quantifies both the number of observed OTUs and their dominance in the community (see Supplementary Information Sec. 5.1.1 and 5.1.2).

We first compare the cohorts from each of the projects, HMP and SMP, separately. Figure 4b shows that human-associated microbial communities from different body sites exhibit a pattern of constrained complexity, i.e., communities with a large number of species tend to have low effective connectance, and vice-versa. A similar pattern is observed when analysing the sponge-associated microbial communities from different geographical locations (Fig. 4c). Notably, when all the cohorts from both studies are plotted together (Fig. 4d), they mutually exhibit the same qualitative pattern. Moreover, the communities of both studies do not represent distinct sizes but rather a pronounced number of them are in an overlapping *n*_eff_ range and are well described by a single stability-complexity empirical upper limit. This may suggest universal, host-independent stability constraints for microbial ecosystems.

The statistical significance of the patterns observed in Fig. 4a-d is assessed by finding a curve of the form *D*^2^ = *βn^−γ^* (*β* and *γ* are fitting parameters) that characterizes the empirical upper limit of the data points (see Methods and Supplementary Information Sec. 10). The associated one-tailed P values of these curves, calculated by comparison to shuffled data realizations in which there is no stability-complexity relationship, are *P* = 1.1 × 10^−2^, *P* = 10^−3^, *P* = 2.7 × 10^−2^, and *P* < 10^−3^. In addition, the Pearson correlation coefficients are *ρ* = −0.528, *ρ* = −0.75, *ρ* = −0.51 and *ρ* = −0.52, and the associated P values are *P* = 0.016, *P* = 0.003, *P* = 0.05, and *P* = 0.003, respectively. These observations strongly suggest the existence of a stability-complexity relationship as first predicted by May, in natural microbial ecosystems.

To test whether our results could be merely an outcome of the differences between the abundance distributions of the species in the different communities, rather than the interrelations between them, we repeat the analysis on shuffled OTU tables. These shuffled tables preserve the abundance distribution of each OTU but eliminate the effect of inter-species interactions. We have established that the measured DOC was effectively flat in all cases, and the complexity-stability pattern could not be reproduced (See Supplementary Information Sec. 7).

The observed patterns could have alternatively been explained as a result of a bias due to a sampling process from a shared species pool, without any inter-species interactions. Communities with small numbers of species are more likely to ‘mirror’ the abundance distribution of the pool, an effect that leads to a steeper DOC (See Supplementary Information Sec. 8). However, we find that this model fails to describe real metagenomic data, as the samples with high overlap all mimic the pool and subsequently each other, while in real data there are clear signs of multiple steady states, so that the high-overlap samples do not all fall into a single shared composition (see Supplementary Fig. 15).

Finally, the advantage of using a top-down methodology is apparent, as the complexity-stability pattern in microbial communities is not observed when the connectivity is extracted from correlation networks (see Supplementary Information Sec. 9).

## Discussion

The governing mechanisms that determine the species’ richness in microbial ecosystem are varied, including assembly processes, such as dispersal limit or habitat filtering, and dynamic processes due to a tangled network of interspecific interactions [43–46]. The observed trade-off between the number of species and the effective connectance emphasizes the role of the later. Accordingly, in order to maintain stability, natural microbial ecosystems had to evolve to either small and highly interacting, or to large but weakly interacting communities.

Our methodology for evaluating the effective connectance of ecosystems can also be applied to other microbial communities from other environments, as well as to general ecological communities. It can also be useful beyond studying the complexity-stability relationship. Even when a system is stable, the effective connectance may determine the impact of local perturbations on the entire system. For example, a high effective connectance suggests that a local perturbation in the abundance of one or a few species is expected to propagate and affect the entire community much more significantly than a system with low connectance.

## Acknowledgements

A.B. thanks the Azrieli Foundation for supporting this research.

## Conflict of Interest

The authors declare no conflict of interest.

## Methods

### Calculating the slope of the DOC

Consider two microbial samples, represented by two abundance vectors of *N* species, **x** = (*x*_1_*,…, x_N_*) ∈ ℝ^*N*^ and **y** = (*y*_1_*,…, y_N_*) ∈ ℝ^*N*^, where *x_i_* and *y_i_* are the relative abundances of species *i* in the two samples, respectively, so that 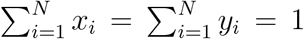. The sets of present species (species assemblages) of **x** and **y** are defined as *X* = {*i*|*x_i_* > 0} and *Y* = {*i, y_i_* > 0}. The overlap between the two samples is given by:

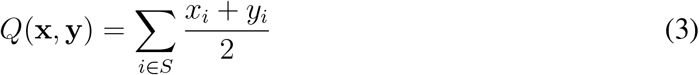

where *S* ≡ *X* ∩ *Y* is the set of shared species present in both samples. If S is empty, *Q*(**x**, **y**) = 0. If *S* = {1*,…, N*}, that is, all the species in *X* and *Y* are shared, then *Q*(**x**, **y**) = 1, but the abundances of **x** and **y** can still differ.

To compare the abundance profiles of two samples, we first renormalize the relative abundances of only the shared species (in set *S*), yielding 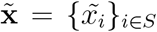 and 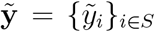. Here, 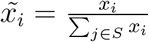, and 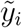 is defined similarly. Then we calculate the dissimilarity between 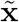 and 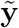. In this paper, we used two dissimilarity functions: i) The rJSD is calculated as:

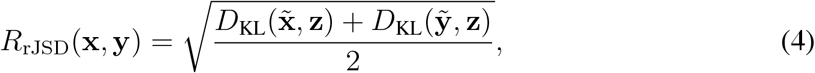

in which 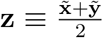, and 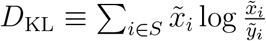 is the Kulleback-Leibler diverrgence between 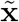 and 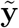. ii) The Euclidean distance between 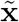 and 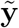 is defined as:

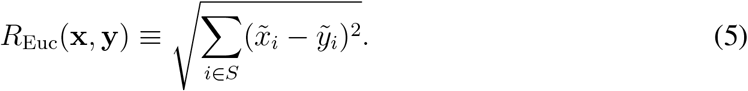

To calculate the DOC’s slope, *D*, in a cohort of *m* samples, we first calculate the overlap *Q_j_* and dissimilarity *R_j_* between all sample pairs, *j* = {1*,…, m*(*m* − 1)/2}. Then, *D* is calculated as the slope of the linear regression model *R_j_* = *b*_0_ + *DQ_j_* + *ϵ_j_*,describing the dissimilarity-overlap relation between the 25% of the pairs with the highest overlap values.

### Metagenomic data sets used in this work

To analyse the complexity-stability patterns of real microbial communities, we used two large-scale cross-sectional microbiome data sets. (1) The Human Microbiome Project (HMP)[12], a 16S rRNA gene-based data set of the human microbiomes from 239 healthy subjects. After the pre-processing and filtering steps detailed below (see ‘Pre-processing steps’ section), we analyzed 13 body sites in four areas: the oral cavity (nine sites: saliva (m=163), tongue dorsum (m=184), palatine tonsils (m=181), keratinized gingiva (m=185), hard palate (m=172), buccal mucosa (m=179), throat (m=172), and sub- and supragingival plaques (m=179 and m=183, respectively)), the gut (one site: stool (m=184)), the nasal cavity (one site: anterior nares (m=145)), and the skin (two sites: left and right retroauricular crease (m=159 and m=163, respectively). Full protocol details are available at the HMP DACC website (http://hmpdacc.org/HMMCP). We performed the DOC analysis at the OTU level. We used a single sample from each subject. In cases where more than one sample was available, we used the first collection. (2) The Sponge Microbiome Project (SMP) [13], a 16S rRNA gene-based data set of the sponge microbiomes from sponges at different geographical locations. After the pre-processing steps detailed below, the data set analyzed includes 15 cohorts of samples from 6 countries: Australia (two cohorts, m=[88, 76]), the Commonwealth of Puerto Rico (one cohort, m=35), Guam (one cohort, m=54), Norway (one cohort, m=73), Spain (six cohorts, m=[80, 65, 44, 70, 41, 41]), and the USA (four cohorts, m=[59, 60, 33, 57]).

### Pre-processing

#### Defining cohorts of SMP samples

The SMP data includes 3,569 microbial samples from diverse geographical locations. To ensure homogeneous cohorts, which are needed for the DOC analysis, we first divide the samples into their respective countries of origin. Next, we remove from the samples of each country all species that are not present in any of the samples. Then, we calculate the Jaccard distances between all sample pairs of each country, based on the presence/absence of the species. The Jaccard distance matrix is then used to calculate and plot the *t*-SNE and PCoA plots of reduced dimensionality, as well as the histogram of the between-samples distances. Through visual inspection of the above mentioned plots, we estimate the number of appropriate clusters for each country, and divide the samples using a k-medoids clustering algorithm (`kmedoids` function in MATLAB) with 10 replicates, based on the matrices of between-samples Jaccard distance.

#### Filtering outliers and low-abundance OTUs

The cohorts of samples from both the HMP and the SMP are pre-proccessed as follows. First, we remove from each cohort low-abundance OTUs with less than 1 read per sample on average, across all the samples in the cohort. To identify and remove ‘outliers’ samples, we calculate the Jaccard distance of each sample with all the other samples, and calculate for each sample the average distance to the other samples. Then, we find a ‘representative sample’, *m′*, whose mean Jaccard distance to all other samples is minimal. We remove all the samples with a Jaccard distance from *m′* greater than two standard deviations from the mean Jaccard distance from *m′*. We filter out any remaining clusters with less then 35 samples, to allow estimation of the DOC’s slope (see Supplementary Information Sec. 4). Lastly, since the DOC’s slope, *D*, is defined only at the high overlap region, we calculate the overlap *Q* between all sample pairs in the cohort, and include in our analysis only cohorts with mean overlaps larger than 0.5.

### Statistical test for the complexity-stability patterns

To calculate the statistical significance of the observed patterns of *D*^2^ versus *n* (Fig. 4), we perform two independent tests: 1) To asses the general trend of a negative relation between *D*^2^ and *n*, we calculate the Pearson correlation and the associated P value; 2) To asses the constrained complexity of the points of *D*^2^-*n* pairs, we find a curve that represents the empirical upper boundary of the points, and calculate its statistical significance with respect to randomized realizations, as follows. We find a curve of the form *g*(*x*) = *βx^−γ^*, where *β* and *γ* are fitting parameters, such that at least 90% of the points are below the curve 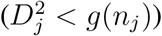, with the minimal ratio *r* between the number of data points and the number of possible *D*^2^-*n* combinations below it. Specifically, the possible *D*^2^-*n* combinations are defined as all possible pairs of *D*^2^ and *n* selected independently from the observed values. The ratio *r* was defined to be the test statistic. We follow the same procedure of finding the optimal curve, and calculating the associated test statistic *r* for 10^3^ Monte-Carlo realizations of shuffled data, i.e., the *n* values are randomly assigned to the *D*^2^ values. The P value is calculated as the fraction of randomized realizations, with *r* equal or larger than the real data. For a detailed description of the optimization procedure see Supplementary Information Sec. 10.

### Population dynamics model

The Generalized Lotka Volterra (GLV) model represents the dynamics of *N* interacting species as a set of ordinary differential equations,

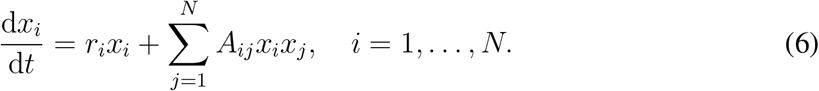

In our simulations, the growth rate of each species *r_i_* is randomly drawn from a uniform distribution 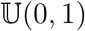, the inter-species interactions coefficients *A_ij_* are randomly chosen from a normal distribution ℕ(0*, σ*^2^) with probability *C*, and are 0 otherwise. The intraspecific coefficients, *A_ii_*, are set to −1 for all species. For a particular GLV model, a cohort of ‘samples’ is simulated as a set of alternative steady states. Different steady states are generated by choosing different initial conditions **x**(*t* = 0). The species assemblage of each sample *ν* is randomly chosen, where each species is present with probability *p^ν^*, which is randomly chosen from a uniform distribution 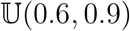. The initial abundances of the present species are subsequently chosen from a uniform distribution 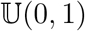. The steady state is simulated by integrating the GLV differential equation using the `ode45` function in MATLAB.

### Demonstration of a complexity-stability relationship in GLV modelsa

In order to generate GLV models with constrained complexity due to stability requirements, we randomly choose pairs of *C* and *σ*, and find the GLV model with the largest number of species that satisfies the system’s stability, i.e., the GLV differential equations can be integrated to a steady state that includes all species. Then, we generate a cohort of *m* = 50 different steady states from each GLV model. We repeat this process to simulate 20 different cohorts defined by different pairs of *C* and *σ*. For each cohort, we apply the DOC analysis and evaluate *D*^2^. The cohorts with high effective connectance (high *D*^2^) can be stable only when the average number of species *n* is relatively small, while large numbers of species are found only in cohorts with low effective connectance, as expected from May’s stability criterion. Importantly, when Euclidean distance is used to calculate the DOC, the data points of the GLV simulations are indeed closer to the predicted stability limit (See Supplementary Information Sec. 5.2.1).

## Notes

### Competing Interest Statement

The authors have declared no competing interest.

### Summary of Updates

One author's name was corrected, the authors' order was changed, and the ORCID numbers were added.

